# Natural Selection Constrains Neutral Diversity Across a Wide Range of Species

**DOI:** 10.1101/006122

**Authors:** Russell B. Corbett-Detig, Daniel L. Hartl, Timothy B. Sackton

## Abstract

The neutral theory of molecular evolution predicts that the amount of neutral polymorphisms within a species will increase proportionally with the census population size (Nc). However, this prediction has not been borne out in practice: while the range of Nc spans many orders of magnitude, levels of genetic diversity within species fall in a comparatively narrow range. Although theoretical arguments have invoked the increased efficacy of natural selection in larger populations to explain this discrepancy, few direct empirical tests of this hypothesis have been conducted. In this work, we provide a direct test of this hypothesis using population genomic data from a wide range of taxonomically diverse species. To do this, we relied on the fact that the impact of natural selection on linked neutral diversity depends on the local recombinational environment. In regions of relatively low recombination, selected variants affect more neutral sites through linkage, and the resulting correlation between recombination and polymorphism allows a quantitative assessment of the magnitude of the impact of selection on linked neutral diversity. By comparing whole-genome polymorphism data and genetic maps using a coalescent modeling framework, we estimate the degree to which natural selection reduces linked neutral diversity for 40 species of obligately sexual eukaryotes. We then show that the magnitude of the impact of natural selection is positively correlated with Nc, based on body size and species range as proxies for census population size. These results demonstrate that natural selection removes more variation at linked neutral sites in species with large Nc than those with small Nc, and provides direct empirical evidence that natural selection constrains levels of neutral genetic diversity across many species. This implies that natural selection may provide an explanation for this longstanding paradox of population genetics.

## Introduction

The level of neutral genetic diversity within populations is a central parameter for understanding the demographic histories of populations [1], selective constraints [2], the molecular basis of adaptive evolution [3], genome-wide associations with disease [4], and conservation genetics [5]. Consequentially, numerous empirical surveys have sought to quantify the levels of neutral nucleotide diversity within species, and considerable theory has focused on understanding and predicting the distribution of genetic variation among species. All else being equal, under simple neutral models of evolution, levels of neutral genetic diversity within species are expected to increase proportionally with the number of breeding individuals (the census population size, Nc). Although this prediction is firmly established, surveys of levels of genetic variation across species have revealed little or no correlation between levels of genetic diversity and population size [6–9]. This discrepancy—first pointed out by Richard Lewontin in 1974 [6]—remains among the longest standing paradoxes of population genetics.

One possible explanation for this disagreement is an inverse correlation between mutation rate and population size. This is expected if there is relatively weak selection against alleles that cause higher mutation rates [8, 10]. Alternatively, this paradox could result from greater impact in large populations of non-equilibrium demographic perturbations such as higher variance in reproductive success [11] or population-size fluctuations [12]. Indeed one recent empirical study suggests that demographic factors play an important role in shaping levels of genetic diversity within animal populations [13]. However, none of these potential explanations is sufficient to fully account for the observed patterns of neutral diversity across species [8].

Another potential cause of this paradox is the operation of natural selection on the genome [7,14,15]. Natural selection can impact levels of neutral diversity via the adaptive fixation of beneficial mutations (hitch-hiking) [7,15,16] and/or selection against deleterious mutations (background selection) [17,18]. Both processes purge neutral variants that are linked to selected mutations, implying that if natural selection is sufficiently common in the genome it can reduce observed levels of neutral polymorphism. Furthermore, theoretical arguments [7,14,13] suggest that, when the impact of natural selection is substantial, the dependence of neutral diversity on population size is weak or even nonexistent. Although many authors have demonstrated that natural selection could, in principle, be sufficiently common to explain Lewontin’s paradox [7,8,14–16,20], few direct empirical tests of this explanation exist.

One unique prediction of the hypothesis that natural selection is a primary contributor to disparity between Nc and levels of neutral genetic variation within species is that natural selection will play a greater role in shaping the distribution of neutral genetic variation in species with large Nc. To test this prediction, we relied on the fact that the impact of natural selection on linked neutral diversity depends on the local recombinational environment. In regions of relatively low recombination, selected variants affect more neutral sites through linkage, and vice versa in regions of relatively high recombination. The resulting correlation between recombination and polymorphism [21–26] (reviewed in depth in [27]) allows a quantitative assessment of the magnitude of the impact of selection on linked neutral diversity (e.g. [22,23,26,28]). Specifically, if the effects of linked selection can explain the lack of correlation between neutral diversity and population size, we expect that species with larger population sizes will display stronger correlations between recombination and polymorphism than those with smaller population sizes, and show a concurrently larger impact of natural selection on levels of neutral diversity across the genome.

Although empirical studies that explore the relationship between neutral diversity and population size are relatively infrequent compared to theoretical studies on this topic, two interesting patterns that merit consideration here. First, the proportion of non-synonymous substitutions that have been driven to fixation by positive selection varies widely across taxa. In humans [29], yeast [30], and many plant species [31], estimates of this proportion are close to zero. In contrast, in *Drosophila* [32,33], mice [34], *Capsella grandiflora* [35], as well as other taxa (reviewed in [8]), a large fraction of non-synonymous substitutions are inferred to have been driven to fixation by positive selection, implying that natural selection is common in the genomes of these organisms (which generally have large Nc). Second, the strength of the correlation between polymorphism and recombination varies widely among the limited number of taxa [8,27] which have been studied in depth. Here again, *Drosophila* [21,25,36] is among the taxa that shows the strongest correlation and thus the clearest evidence for natural selection, and the correlation in *Drosophila* is substantially larger than, for example, humans [28].

In a related study to the work presented here, Bazin *et al.* [37] showed that there is no correlation between nucleotide diversity in non-recombining mtDNA and nucleotide diversity in the nuclear genome. While this is consistent with some predictions of theoretical work on this subject, the mitochondrion has unusual patterns of replication and inheritance and it is therefore challenging to disentangle the processes that generate diversity from those that shape its distribution across the genome. Although suggestive, the evidence accrued thus far is fragmentary, has not been analyzed in aggregate, and varies widely in quality of samples, data collection, and analyses performed [8,27]. It is therefore difficult to draw firm conclusions about the relative importance and prevalence of natural selection in shaping patterns of genetic variation in the genome based on existing studies.

Due to rapid advances in genome sequencing technologies, whole-genome polymorphism data are now available for a wide variety of species (e.g. [36,38]), and these data enable us to conduct a quantitative test of the natural selection hypothesis as an explanation for Lewontin’s paradox. Towards this, we identified 40 species with sufficiently high quality reference genomes, linkage maps, and polymorphism data to enable a broad-scale, robust comparison of the relative strength of correlation between polymorphism and recombination rate within a single unified alignment, assembly, and analysis framework. Using these data, and reasonable proxies for Nc, we show that the effect of selection on linked nucleotide diversity is indeed strongly correlated population size. In other words, natural selection plays a disproportionately large role in shaping patterns of genetic variation in species with large Nc, confirming the idea that natural selection is an important contributor to Lewontin’s paradox.

## Results

### Genomic datasets and modeling approach

We identified 40 species (15 plants, 6 insects, 2 nematodes, 3 birds, 5 fishes, and 9 mammals) for which a high quality reference genome, a high density, pedigree-based linkage map, and genome-wide resequencing data from at least two unrelated chromosomes within a population were available (Table 1) and Supplemental Tables S1 and S2). Because our model (below) requires that recombination has been sufficiently frequent to uncouple genealogies across large tracts of DNA on chromosomes, we required that each species have an obligatory sexual portion of its life cycle. This requirement necessarily excludes clades such as bacteria, which are predominantly clonally propagated.

**Table 1:** List of species used in this work

| Species | Common Name | Kingdom | Subgroup |
| --- | --- | --- | --- |
| *Anopheles gambiae* | African malaria mosquito | Animalia | Invertebrate |
| *Apis mellifera scutellata* | Honeybee | Animalia | Invertebrate |
| *Arabidopsis thaliana* | Thale cress | Plantae | Herbaceous |
| *Bombyx mandarina* | Silkworm | Animalia | Invertebrate |
| *Bos taurus* | Cow | Animalia | Vertebrate |
| *Brachypodium distachyon* | Purple false brome | Plantae | Herbaceous |
| *Caenorhabditis briggsae* | Roundworm | Animalia | Invertebrate |
| *Caenorhabditis elegans* | Roundworm | Animalia | Invertebrate |
| *Canis lupus* | Wolf | Animalia | Vertebrate |
| *Capsella rubella* | Pink Shepherd's Purse | Plantae | Herbaceous |
| *Citrullus lanatus lanatus* | Watermellon | Plantae | Herbaceous |
| *Citrus reticulata* | Mandarin Orange | Plantae | Woody |
| *Cucumis sativus* var. *hardwickii* | Cucumber | Plantae | Herbaceous |
| *Cynoglossus semilaevis* | Tongue sole | Animalia | Vertebrate |
| *Danio rerio* | Zebrafish | Animalia | Vertebrate |
| *Drosophila melanogaster* | Fruitfly | Animalia | Invertebrate |
| *Drosophila pseudoobscura* | Fruitfly | Animalia | Invertebrate |
| *Equus ferus przewalskii* | Prewalskii's horse | Animalia | Vertebrate |
| *Ficedula albicollis* | Collared flycatcher | Animalia | Vertebrate |
| *Gallus gallus* | Chicken | Animalia | Vertebrate |
| *Gasterosteus aculeatus* | Stickleback | Animalia | Vertebrate |
| *Glycine soja* | Wild soybean | Plantae | Herbaceous |
| *Gossypium raimondii* | New world cotton | Plantae | Woody |
| *Heliconius melpomene melpomene* | Postman butterfly | Animalia | Invertebrate |
| *Homo sapiens* | Human | Animalia | Vertebrate |
| *Lepisosteus oculatus* | Spotted gar | Animalia | Vertebrate |
| *Macaca mulatta* | Rhesus macaque | Animalia | Vertebrate |
| *Medicago truncatula* | Barrel medic | Plantae | Herbaceous |
| *Meleagris gallopavo* | Turkey | Animalia | Vertebrate |
| *Mus musculus castaneus* | House mouse | Animalia | Vertebrate |
| *Oryza rufipogon* | Wild rice | Plantae | Herbaceous |
| *Oryzias latipes* | Medaka | Animalia | Vertebrate |
| *Ovis canadensis* | Bighorn sheep | Animalia | Vertebrate |
| *Papio anubis* | Olive baboon | Animalia | Vertebrate |
| *Populus trichocarpa* | Black cottonwood | Plantae | Woody |
| *Prunus davidiana* | David's peach | Plantae | Woody |
| *Setaria italica* | Foxtail millet | Plantae | Herbaceous |
| *Sorghum bicolor* ssp. *verticilliflorum* | Wild Sudan grass | Plantae | Herbaceous |
| *Sus scrofa* | Wild boar | Animalia | Vertebrate |
| *Zea mays* ssp. *parviglumis* | Teosinte | Plantae | Herbaceous |

Nonetheless, extending this framework to bacterial taxa will be an important step towards understanding the mechanisms by which natural selection shapes patterns of variation across the tree of life. Additionally, our sampling is biased towards more commonly studied clades (e.g., mammals), but this is unavoidable in this type of analysis, and there is no reason in principle why this taxonomic bias would affect the basic conclusions we describe here, as the sampled taxa likely span a large range of census population sizes.

After acquiring sequence data, we developed and implemented a bioinformatic pipeline to align, curate, and call genotype data for each species (see Figure S1 and methods for a full description of the bioinformatics pipeline). We further used the available genetic maps to estimate recombination rates across the genomes. Across all species, we analyzed recombination across nearly 385,000 markers and aligned more than 63 billion short reads. This is therefore the largest comparative population genomics dataset that has been assembled to date.

We used both simple nonparametric correlations, and explicit coalescent models, to test for a relationship between the impact of selection on linked neutral diversity and census size. Although correlations between recombination rate and neutral diversity are informative, the extensive literature in theoretical population genetics provides an opportunity to develop a robust modeling approach. Two primary types of selection can introduce a correlation between recombination rate and levels of nucleotide diversity: background selection (BGS) and hitchhiking (HH). Here, we are not primarily largely concerned with distinguishing between the two models, and so focus on their joint effects. In addition to combining background selection and hitchhiking, we would also like to relax the assumption that these processes act uniformly across the genome. All else being equal, regions of the genome with a higher density of potential targets of selection should experience a greater reduction in neutral diversity.

Starting from considerable prior theoretical work [14,17,18,32,39–41], we develop an explicit model relating polymorphism, recombination rate, and density of functional elements in the genome. We fit both a joint model that allows for both HH and BGS, as well as models of BGS only, HH only, and a purely neutral model (in which there is no predicted correlation between recombination or functional density and neutral diversity). Using these models, we estimate the fraction of neutral diversity removed by linked selection for beneficial alleles and/or against deleterious alleles (Figure 1) for each species, as well as the relative likelihood of each model.

**Figure 1.**
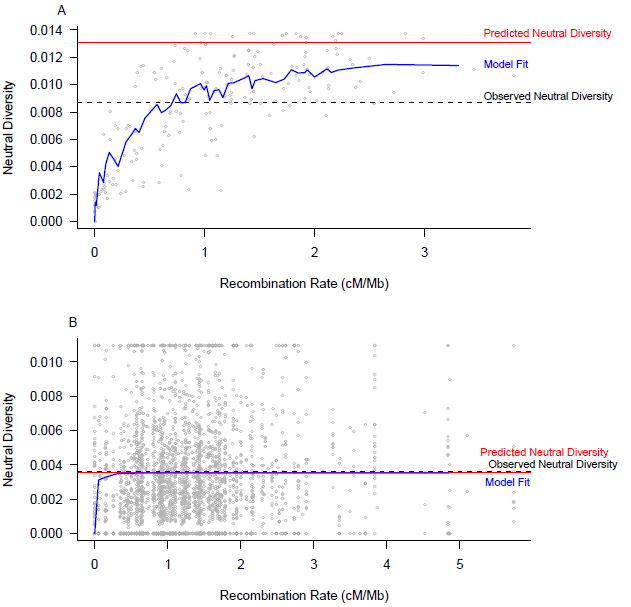
Estimating the impact of selection on linked neutral variation. To obtain a direct estimate of the amount of linked neutral variation removed by selection, we fit a population genetic model incorporating hitchhiking and background selection effects to the estimates of pi and recombination rate in 500 kb windows across the genome. Model fit (blue), estimated neutral diversity in the absence of selection (red), and observed genetic diversity (dashed) are shown for a species with large population size (*D. melanogaster*, part A) and small population size (*Equus ferusprzewalskii,* part B). The magnitude of the impact of selection on linked neutral diversity is estimated as 1 – (observed neutral diversity / neutral diversity in the absence of selection).

In practice it is not feasible to determine Nc for the majority of species we studied. Instead, we used the species’ geographic range and individual body size as proxies for Nc. Size has been previously validated as a proxy for individual density in a wide variety of taxa and ecosystems (e.g. [42–44]). Under some simplifying assumptions, the product of geographic range and local density should be sufficient to roughly estimate a species census population size, and each factor is expected to independently capture some information related to species’ Nc. Specifically, we expect that range will be positively correlated with Nc, size will be negatively correlated with Nc, and Nc will be positively correlated with the impact of selection.

### Natural selection removes more linked neutral variation in species with large census sizes

For many of the species that we studied, it is clear that selection plays a central, even dominant, role in shaping patterns of neutral genetic diversity. Specifically, both our correlation analysis and our explicit modeling support the hypothesis that natural selection on linked sites eliminates disproportionately more neutral polymorphism in species with large Nc, and in this way natural selection truncates the distribution of neutral genetic diversity.

At a coarse scale, there is a stronger correlation between polymorphism and recombination in invertebrates (mean partial τ after correcting for gene density = 00.247), which likely have a large Nc on average, than in vertebrates (median partial τ = 0.118), which likely have smaller Nc on average (twotailed permutation P = 0.021). We observe similar patterns for herbaceous plants (mean partial τ = 0.106) versus woody plants (mean partial τ = −0.020; two-tailed permutation P = 0.058) and for medians as opposed to means (Figure 2). When we repeat the analysis with alternate window sizes, we observe consistent effect sizes, albeit occasionally with reduced statistical support (Supplemental Table S3).

**Figure 2.**
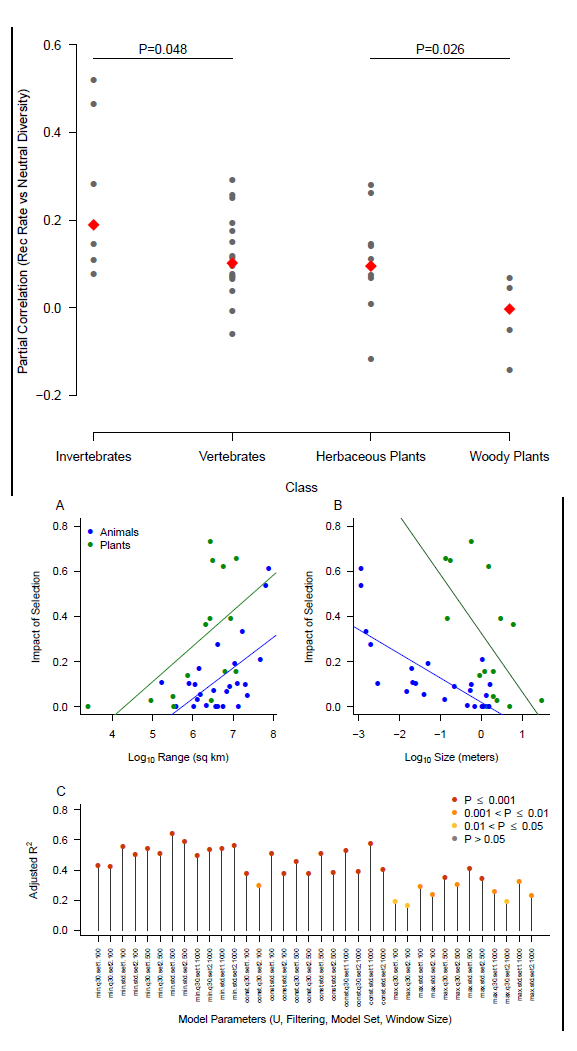
The correlation between neutral diversity and recombination rate is stronger in taxonomic classes expected to have larger population sizes. We estimated both neutral diversity and recombination rate in 500-kb windows across the genome and computed partial correlations (Kendall’s τ after accounting for variation in functional density (measured as proportion of sites in a window that are part of an annotated protein-coding exon). The significance of differences in median τ (red diamonds) between vertebrates and invertebrates, or between woody and herbaceous plants, is based on Wilcoxon Rank Sum Tests (***P<0.001, ** P<0.01, * P<0.05).

More generally, we tested the hypothesis that Nc is positively correlated with the impact of selection by fitting a linear model that includes body size, geographic range, kingdom and the significant interactions among them as predictors, and uses the impact of selection estimated from our coalescent model as the response variable (Table 2; Figure 3)Table Coding). Both size and range are significant predictors of the impact of selection in the expected directions (Table 2; log10(size): coefficient = −0.092, P = 0.0005; log10(range): coefficient = 0.112, P = 0.0002), and model as a whole explains 63.88% of the variation in impact of selection across species (Table 2; overall P = 3.518 × 10^−8^). This is clear evidence that more variation is removed from the genomes of species with smaller body size and larger ranges than from the genomes of species with larger body size and smaller ranges.

**Figure 3.**
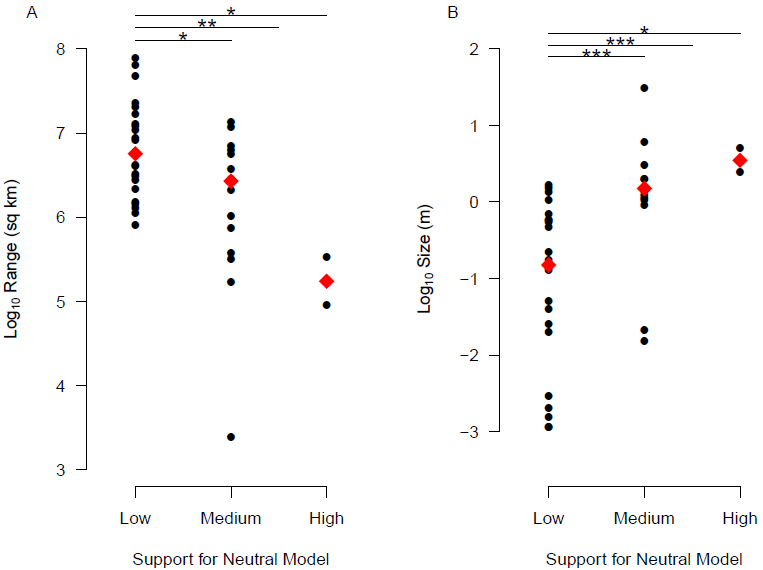
Proxies for census size are correlated with the estimated impact of selection on neutral diversity. For each species, we obtained estimates of size (in meters) and range (in sqaure kilometers), and used those as predictors in a linear model with a measure of the impact of selection on neutral diversity as the response (see main text and Table 2 for full model information). Both size (part A) and range (part B) are significant predictors of the impact of selection on neutral diversity, in the expected directions. Points are colored by kingdom (blue=animals, green=plants), and regression lines estimated independently for plants and animals are shown. C) Robustness of our model fit. We tested our main model (see text and Table 2) across a wide range of different analysis options, including different SNP filtering options, different window sizes, and different population genetic parameters. Each point represents the adjusted R^2^ of the full model for one set of parameter values, colored according to P-value.

**Table 2:** Linear model fit for the main model

|  | Estimate | Std. Error | t value | Pr(>\|t\|) |
| --- | --- | --- | --- | --- |
| (Intercept) | -0.71408 | 0.18282 | -3.906 | 0.000410 |
| Log<sub>10</sub> (range) | 0.11239 | 0.02742 | 4.100 | 0.000234 |
| Log<sub>10</sub> (size) | -0.09229 | 0.02414 | -3.823 | 0.000520 |
| Kingdom (0=animal, 1=plant) | 0.34567 | 0.05091 | 6.789 | 7.16e-08 |
| Log<sub>10</sub> (size) : Kingdom | -0.12337 | 0.06075 | -2.031 | 0.049935 |

2 Overall F-statistic: 18.24 on 4 and 35 DF, p-value: 3.518e-08, adjusted R-squared: 0.6388

A number of confounding factors could potentially influence our conclusions, including variation in map or assembly quality across species, differences in overall recombination rate, and differences in genome size. To test whether these factors can explain our results, we fit a confounder-only model including two measures of genetic map quality (density of useable markers and proportion of total markers scored as useable); two measures of assembly quality (proportion of assembly that is not gaps and proportion of total assembly assembled into chromosomes); overall recombination rate; and genome size. We then compare this confounder-only model to a model that includes all confounding parameters, and in addition includes our population size proxies (kingdom, size, range). The model with proxies for Nc both explains substantially more total variation in impact of selection (adjusted R^2^ 0.6359 compared to 0.3388 for the confounder-only model) and is a significantly better fit to the data (F=7.7322, df=4, P=0.0002).

To get a lower bound on the proportion of variation in impact of selection explained by our parameters of interest (range, size, kingdom, and the kingdom*size interaction), we fit a linear model with these parameters as predictors and the residuals of the confounder-only model as the response variable (Supplemental Tables S4, S5). This is a conservative test, as genome size is strongly correlated with body size (Kendall’s τ = 0.296, P=0.007 in our dataset). Nonetheless, our proxies for Nc explain 34.05% of the remaining variation in impact of selection after accounting for all confounding parameters (overall model P = 0.0008, Table S4), and 47.36% of the variation after accounting for all confounding parameters except genome size (overall model P = 2.042 × 10^−5^, Table S5).

For five species, our polymorphism data included individuals from domesticated populations, which could potentially affect our conclusions if selection has a different signature during domestication events than it leaves in natural populations. However, removing these five species has virtually no impact on our model fit (overall adjusted R^2^ = 0.6281, overall P = 6.094 × 10^−7^, Supplemental Table S6), suggesting that their inclusion has not biased our results. Additionally, we obtain similar results if we fit our model (excluding the kingdom term) to animals and plants independently (Supplemental Table S7 and S8). Finally, varying the SNP filtering criteria, window size, assumed deleterious mutation rate (U), or population genetic modeling approach produces nearly identical results (Figure 3C), implying our primary conclusion is robust to a wide range of analysis choices. Taken together, our analysis demonstrates that the central pattern - natural selection reduces neutral diversity more strongly in species with large Nc than in species with small Nc - is consistently observed with both nonparametric model free approaches (Figure 2; Supplemental Table S3) and with explicit population genetic models (Figure 3A,B, (Table 2)) across a wide range of possible analyis and filtering choices (Figure 3B, Supplemental Tables S4-S8).

If the process of recombination is itself mutagenic, neutral processes could produce a correlation between recombination and polymorphism [21,25,27]. However, no or very weak correlations between divergence and recombination has been found in most species that have been closely studied [21,25] (reviewed in [27]). Moreover, for those species in which a positive correlation between divergence and polymorphism has been found *(e.g.* [45,46]), it is likely at least partially the result of linked selection acting on polymorphisms present in the ancestral population [27,32]. Furthermore, the two species that showed the strongest correlation between polymorphism and recombination (partial τ = 0.5196 for *D. melanogaster*, partial τ = 0.4637 for *D. pseudoobscura*) have no such correlation between recombination rate and divergence either on broad scales [21] or fine scales [25]. Moreover, many authors have found strong evidence that recombination is not mutagenic in a number of other animal species (e.g. [28,47,48]), and it therefore appears a general consensus has emerged that recombination-associated mutagenesis is unlikely to influence the overall patterns we report in this work [27].

### Species with small census sizes show stronger evidence for neutrality

As an alternative approach to estimating the impact of natural selection on linked neutral diversity, we considered whether our proxies for Nc correlates with the strength of evidence that selection shapes patterns of neutral diversity, derived from our population genetic modeling approach. To do this, we focus on the relative likelihoods (Akaike weights) of four models: the BGS+HH model, the BGS-only model, the HH-only model, and the neutral model. These relative likelihoods can be interpreted as the probability that a particular model is the best model according to AIC, given the set of models tested and the underlying data.

We initially focus on the relative likelihood of the support for a purely neutral model. Species with weak or no support for neutrality (relative likelihood of the neutral model < 0.05) have significantly larger ranges (P=0.006, Wilcoxon Rank Sum Test, (Figure 4A) and significantly smaller sizes (P=0.0001, Wilcoxon Rank Sum Test, (Figure 4B) than species with moderate (relative likelihood of neutral model > = 0.05 and <0.90) or strong (relative likelihood of neutral model > =0.90) support. This pattern also holds if we compare the species with strong support for neutrality or species with moderate support for neutrality individually to species with weak or no support (moderate vs weak: P= 0.0005 for size and 0.02 for range; strong vs weak: P=0.02 for size and 0.02 for range, all P-values from Wilcoxcon Rank Sum Tests). This suggests that the evidence for non-neutral processes (background selection and/or hitchhiking) is significantly stronger in species with larger ranges and/or smaller sizes, consistent with our results above and with the hypothesis that natural selection explains Lewontin’s paradox.

**Figure 4.**
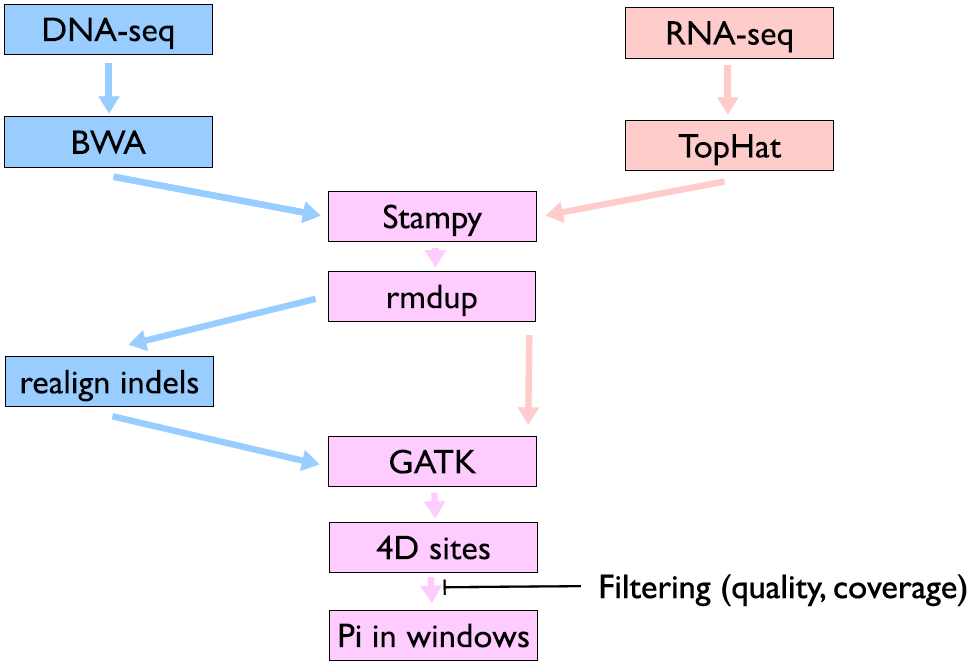
Species with little evidence for selection have smaller census sizes. For each species, we estimated the relative likelihood of a purely neutral model (no impact of selection on neutral diversity), based on AIC values for neutral and selection models (see methods for details), and categorized species based on support for neutrality (low=relative likelihood<0.05, medium = relative likelihood > 0.05 but < 0.5, high=relative likelihood>0.50). Species with more support for neutrality have smaller ranges (part A) and larger body size (part B). P-values for comparisons (indicated by lines at the top of each panel) of low vs. high; low vs. medium; and low vs. medium and high combined are based on Wilcoxon Rank Sum tests (*** P < 0.001, ** P < 0.01, * P < 0.05).

### Hitch-hiking appears more prevalent in large Nc species

Given the extensive debate on the relative importance of hitch-hiking versus background selection in shaping patterns of diversity across the genome [17,21], we also attempt to disentangle the relative roles of these two processes in reducing neutral diversity. This is potentially relevant to the resolution of Lewontin’s paradox, as models of frequent, recurring hitch-hiking (i.e., genetic draft [7]) demonstrate that recurrent hitch-hiking can remove the dependence of neutral diversity on population size entirely. Thus, evidence that hitch-hiking specifically is more likely to occur in species with large census sizes would be compelling evidence for a role of selection in resolving the discrepancy between population sizes and neutral diversity. However, it is crucial to note that our test does not take into account features, such as patterns of polymorphism around amino acid fixations [23,49], that are particularly powerful for distinguishing hitch-hiking and background selection, and thus suffers from many of the limitations of previous work relying purely on patterns of neutral diversity across the genome *(e.g.* [26,28,40,41]).

With that caveat, we begin by noting that, consistent with recent work in *Drosophila [49,50]* and other organisms [26,28,48], background selection is ubiquitous. Either the BGS-only model or the BGS+HH model has at least some support (relative likelihood > 0.05) for 95% (38 of 40) of the species we analyzed, and for 90% (36 of 40) of species one of the BGS-containing models was the best fit, as measured by AIC. Thus, it seems clear that, in most cases, background selection is a more appropriate null model for tests of natural selection than strict neutrality.

To test whether species with moderate (relative likelihood of HH or BGS+HH>0.05 and < 0.5) or strong (relative likelihood of HH or BGS+HH>0.5) evidence for hitch-hiking differ from species with little or no evidence for hitch-hiking (relative likelihood of HH or BGS+HH<0.05), we examined our proxies for Nc among these evidence classes. Species with moderate or strong evidence for hitchhiking have significantly larger ranges than species with weak/no evidence for hitch-hiking (P = 0.03, Wilcoxon Rank Sum Test, median range (low)=2681693 sq km, median range (med/high)=5592037 sq km), and these species tend to have smaller sizes as well (P=0.15, Wilcoxon Rank Sum Test, median size (low)=0.91 m, median size (med/high)=0.54 m).

As a secondary test of this pattern, we compared whether the relative likelihood of hitch-hiking was greater for species estimated to have particularly high Nc compared to species estimated to have particularly low Nc. We define the high-Nc class as those species with ranges greater than the median range, and sizes below the median size, and we define the low-Nc class as those species with ranges below the median range and sizes above the median size. The relative likelihood of hitch-hiking models is greater for species in the high-Nc class (P=0.023, Wilcoxon Rank Sum Test), and the proportion of species with moderate or strong evidence for hitch-hiking (either alone or in combination with BGS) is higher in the high-Nc class than the low-Nc class (4/10 in high-Nc class, 0/10 in low-Nc class, P=0.086, Fisher’s Exact Test).

Despite the fact that our test is unlikely to have substantial power to distinguish background selection and hitch-hiking models, we suggest that these results imply that hitch-hiking in particular is a stronger force shaping genomic diversity in species with large Nc, while background selection appears to be much more pervasive. The observation that pervasive hitch-hiking may predominantly occur in species with large Nc suggests that genetic draft may play a substantial role in limiting neutral diversity among the species with the largest population sizes. More data on species with very large Nc, and the application of tests specifically designed to detect hitch-hiking to a wider taxonomic range, will be necessary to fully disentangle the relative roles of hitch-hiking and background selection in shaping levels of neutral diversity.

## Discussion

On the strength of early allozyme polymorphism data, Lewontin [6] observed that in contrast with theoretical predictions of the neutral theory [51–53], the range of neutral genetic variation among species is substantially smaller than the range of Nc among species. Because both positive selection via hitch-hiking and negative selection via background selection purge linked neutral mutations, the operation of natural selection affects patterns of neutral genetic variation at linked sites across the genome. Although many authors have suggested that natural selection may play a role in truncating the distribution of genetic variation and may play a greater role than neutral genetic drift in shaping patterns of neutral nucleotide polymorphism [7,8,14,15], few empirical tests of this hypothesis has been proposed or conducted. Here, we showed that species with larger Nc display a stronger correlation between neutral polymorphism and recombination rate, and that natural selection removes disproportionately more linked neutral variation from species with larger populations. This indicates that natural selection plays a disproportionately large role in shaping patterns of polymorphism in the genome of species with large Nc.

One important consideration when interpreting our results is that cryptic population structure can influence patterns of variation across the genome in a way that obscures the effects of selection. In the extreme case, where populations do not exchange any migrants for an extended period of time, genetic divergence is expected to accumulate at equivalent rates across the genome and would obscure the effects of linked selection. Elucidating the complex relationship between population structure and patterns of natural selection is an important and longstanding question in population genetics (for recent work see *e.g.* [54,55]). Nonetheless, especially given the scope of our analysis, it is not feasible to simultaneously estimate the effects of linked selection and population structure, and there are many reasons to believe that the results presented here will be robust to potential cryptic population structure.

So long as the population subdivision is not especially ancient (in the timescale of coalescence, on the order of Ne generations), a correlation between recombination and polymorphism is expected to remain due to the effects of selection on linked sites in the ancestral population [27,32]. Additionally, if migration is sufficiently common, it is reasonable to treat data derived from samples from separate localities as a single population [56]. One straightforward assumption is that species with larger geographic ranges will have greater opportunity on average to accumulate cryptic population structure than species with small ranges, which would imply we should preferentially underestimate the effects of linked selection in species with larger ranges. If population structure is a primary determinant of patterns of nucleotide diversity in taxa that we studied, we could reasonably expect a negative correlation between species range and the effects of selection on linked sites. Given that we instead obtain the opposite effect—one consistent with the effect of selection on linked neutral sites—it is reasonable to conclude that cryptic population structure has not drastically influenced the basic results presented herein.

Understanding the proximate and ultimate factors that affect the distribution of genetic variation in the genome is a central and enduring goal of population genetics and it carries important implications for a number of evolutionary processes. One implication of this work is that in species with large Nc, such as *D. melanogaster*, selection plays a dominant role in shaping the distribution of molecular variation in the genome. Among other things, this can affect the interpretation of demographic inferences because it indicates that even putatively neutral variants are affected by natural selection at linked sites. Furthermore, to whatever degree standing functional variation is also affected by selection on linked sites (*e.g.* [40]), local recombination rate in organisms with large Nc may also predict what regions of the genome will contribute the greatest adaptive responses when a population is subjected to novel selective pressures.

More broadly, this work provides some of the first direct empirical evidence that the standard neutral theory may be violated across a wide range of species. Indeed, it is clear from this work that in many taxa, natural selection plays a dominant role in shaping patterns of neutral molecular variation in the genome. It is therefore essential that we consider selective processes when we study the distribution of genetic diversity within and between species. Incorporating selection into standard neutral models of evolution will be a central and important challenge for evolutionary geneticists going forward.

## Materials and Methods

### 1. Data sources and curation

Reference genome versions, annotation versions, map references, and other basic information about the genetic and genomic data for species we included in our analysis is summarized in Supplemental Table S1 and S2, and described in more detail below.

### Reference genomes

To identify suitable species for our analysis, we started from the list of genome projects available at GOLD (http://www.genomesonline.org/documents/Export/gold.xls) and NCBI (ftp://ftp.ncbi.nlm.nih.gov/genomes/GENOME_REPORTS/). both accessed 6 October 2013. We removed all non-eukaryotes from both sets. We then further filtered the GOLD set to remove all projects where status was not either “draft” or “complete”, and where project type was not “Whole Genome Sequencing”, and the NCBI set to keep only all projects with MB > 0 and status equal to “scaffold,” “contigs,” or “chromosomes.” Finally, we merged both lists, removed duplicate species, and removed all species without an obligate sexual lifestyle. We required species have an obligatory sexual portion of their lifecycle to ensure that some amount of recombination can be expected in natural populations.

Next, we manually checked the quality of the genome assembly of each species remaining on our list by inspection of assembly reports available from NCBI, Ensembl, Phytozome, or species-specific databases. Any species without chromosome-scale assemblies was removed, as was any species without an available annotation of coding sequence. In two cases (*Heliconius melpomene* and *Gasterosteus aculeatus*), chromosome scale assemblies are available but annotations were only available for the scaffold-level (or a previous, lower quality chromosome-level) assembly. In these cases, we updated the coordinates of the coding sequence annotations using custom Perl scripts (available from the GitHub page associated with this manuscript: see the data accessibility section for details on how to obtain source code and data).

### Polymorphism

We required that each species be represented by random-shearing Illumina short-read sequence data for at least two chromosomes derived from unrelated individuals within the same population. For 4 species (*Bos taurus, Lepisosteus oculatus, Prunus persica, Papio anubis*) we used a single outbred diploid individual. If samples were intentionally inbred or if the species is known to engage in frequent self-fertilization in natural populations, we required data from at least two separate individuals. The number of individuals included and the ploidy of the sequenced individuals are reported in Table S1. For six species, we used polymorphism data from a very closely related taxon to the species that was sequenced to produce the reference genome (Table S1). In particular, we attempted to avoid using polymorphism data from domesticated species where possible; in many cases we were able to use polymorphism data from wild ancestors or close relatives of domesticated plants and animals. Nonetheless, for five species *(Gallus gallus, Bos taurus, Melagris gallopavo, Setaria italica,* and *Citrus reticulata*), we could not identify suitable data from wild populations, and we instead elected to use polymorphism data from heritage breeds and strains (Table S1).

### Genetic maps

We required that each species have available a pedigree-based genetic map, generated from markers that could be mapped to the reference genome by either ePCR or BLAST, and with an average inter-marker spacing (after filtering unmapped and mis-mapped markers; see below) of no more than 10 cM. For species with recombination in both sexes, we used sex-averaged genetic distances where possible, although in two cases (*Anopheles gambiae* and *Bos taurus*) maps were only available for a single sex and so for those species we use a single-sex map by necessity. For species with recombination in only one sex (*e.g. Drosophila*), we corrected genetic distances to represent a sex-averaged value by dividing by dividing estimated recombination rates in the sex with recombination by 2. In 9 cases where genetic maps from the same species as the polymorphism data were unavailable or of insufficient quality (*Zea mays, Prunus persica, Papio anubis, Oryza sativa, Ovis aries, Mus musculus, Equus caballus, Canis lupus familiaris, Citrus clementina) w*e used genetic maps from a closely related taxon, typically the genome species (see Table S1 for full details).

### Range and size information

While ideally we would obtain estimates of actual census population sizes, even moderately accurate estimates are rarely available. As an alternative, we used species range and individual size as proxies for census population size. To determine range, we used occurrence data available from GBIF (http://www.gbif.org/) or published literature (when no occurrence data was available in GBIF) to estimate species distributions as follows. First, for each species, we obtain and then filter all occurrence data stored at GBIF. In general, we filter to require a known source (basis of observation) and exclude fossil records; we also filter to remove clearly erroneous points, such as those well outside the known species range (often arising, e.g., from transposition of longitude and latitude during data entry in museum records) or those falling in oceans for terrestrial organisms and vice versa. Specific filtering steps for each species are documented in the associated R code (available at GitHub). After filtering, we fit an alpha-hull [57] to estimate the species range, which we then filter to remove area overlapping ocean for terrestrial species and overlapping land from oceanic species, and then convert to area by projecting from GPS (WGS84) coordinates to a cylindrical equal area projection using the spTransform function in the R package rgdal (http://cran.r-project.org/web/packages/rgdal/). R scripts to replicate our analysis are provided at the GitHub page associated with this manuscript.

We recognize that GBIF occurrence data reflects current range, and does not account for historical range; however, as accurate long-term historical ranges are not known for most species we are limited to using the data that is available. We also note that for the five domesticated species, plus humans, occurrence data is not of much use for estimating range; in these cases we have attempted to approximate either the historical range of the species (humans, turkey, clementine) or the current range of the heritage breeds the polymorphism sample is obtained from (chicken, cow, millet).

In order to account for variation in population density across species, we also use individual size as a second proxy for Nc. Body size has been validated as a proxy for local population density across a wide variety of systems and taxa (*e.g.* [42–44]), and is readily available from common databases such as Encyclopedia of Life and Animal Diversity Web (Supplemental Table S9). While it would be ideal to obtain quantitative estimates of population densities (*e.g.* by using extensive mark-recapture methods [58]), for the majority of species we studied reliable direct estimates of population-densities are not available.

## 2. Polymorphism pipeline

### Alignment and genotyping pipeline

We acquired short read data from the NCBI short read trace archive. All accession numbers for short read data used in this analysis are listed in Supplemental Table S10. We aligned these data to their respective reference genomes (reference genome versions and relevant citations are listed in Supplemental Table S1). For libraries prepared from genomic DNA we used bwa v0.7.4 [59] wit default options. For libraries prepared from RNA, we aligned reads initially using tophat2 v2.0.7 [60] with default options, except we specified ‘-no-novel-juncs’ and ‘no-coverage-search’ and gave tophat2 a GFF file (version indicated in Table S1) to speed up alignment. For both DNA and RNA we then realigned reads that failed to align confidently using Stampy v1.0.21 [61], with default options. After this, putative PCR duplicates were removed from both RNA and DNA based libraries using the ‘MarkDuplicates’ function in Picard v1.98 (http://picard.sourceforge.net). For DNA libraries, we next use the ‘IndelRealigner’ tool in the GATK v2.4–3 [62] to realign reads surrounding likely indel polymorphisms. These GATK and Picard functions were run using default command line options.

We genotyped all samples using the GATK v2.4–3 [62]. If samples were intentionally inbred, or if the species is known to primarily reproduce through self-fertilization in natural populations we used the ‘-ploidy’ option to set the expected number of chromosomes to 1 (see Supplemental Table S1 for ploidy settings used for each species). We then extracted polymorphism data from four-fold degenerate synonymous sites. While there is mounting evidence that these sites are not evolving under strictly neutral processes (e.g. [63,64]), four-fold degenerate sites are a widely accepted approximation for neutral markers in the genome, and importantly these sites are available in both RNA and DNA sequencing efforts.

We sought to exclude low confidence sites by filtering our genotype data through several basic criteria. First, we required that every fourfold degenerate site have a minimum phred-scaled probability of 20 that there is a segregating site within the sample. To ensure robustness of our results, we also applied a more stringent Q30 genotype quality filter and performed otherwise identical analyses using these data. Second, for every fourfold degenerate site, we computed the mean depth for each sample. We then required each sample have at least half as many reads as the mean depth at a site for that position to be included in the analysis. For variable sites, we further required that phred-scaled strand bias be below 40. This quantity is based on an exact test for how often alternate alleles are called by reads aligned to the > versus the — strand of the reference genome; a large bias might be expected if for example a nearby transposable element insertion relative to the reference genome influenced read alignments on one strand and would make the genotypes at that site unreliable. We further required that the absolute value for the Z-score associated with the read position rank sum, the mapping quality rank sum, and the base quality rank sum be above 4. These statistics quantify how biased the reference allele alignments relative to those of non-reference alleles for the relevant filters. For example, the first filter —read position rank sum—quantifies whether non-reference alleles are generally found further forward or backward in a short read. This filter may also reflect errors due to systematic differences in alignments of non-reference allele bearing reads (*e.g.* due to indels on one of the chromosomes present in an individual). See the GATK [62] documentation for in-depth descriptions of the relevant filters used. We applied these criteria to both DNA and RNA based libraries. Summaries of sites aligned and filtered for each genome are available in Supplemental Table S11, and a schematic of our pipeline is presented in Supplemental Figure S1.

### Homo sapiens

Rather than recompute variant calls, for the human data, we obtained VCF files for the Yoruban population from [38]. We elected to do this because these data are exceedingly well curated and the size of the human variation raw data presents a practical computational challenge. The VCF file was treated as described below in all case.

### Estimating genetic diversity in genomic windows

From these filtered files, we computed genetic diversity as π, the average number of pairwise differences [65], at four-fold degenerate sites in non-overlapping windows of 100kb, 500kb, or 1000kb. In all cases, we excluded windows from our analysis with fewer than 500 sequenced four-fold degenerate sites. We also exclude all windows on sex chromosomes, in order to avoid complicating effects of hemizygosity on patterns of polymorphism.

### 3. Recombination rate estimation pipeline

Our approach to estimating recombination rates is to first obtain sequence information and genetic map positions for markers from the literature, map markers to the genome sequence where necessary, filter duplicate and incongruent markers, and finally estimate recombination rates from the relationship between physical position and genetic position. Specific details of map construction for each species are described in Supplemental Text S1.

### Data curation and mapping markers to the reference genome

We used three basic approaches to link markers from genetic maps to sequence coordinates. In some cases, sequence coordinates are available from the literature, in which case we use previously published values (in some cases updated to the latest version of the genome reference). For cases where primer information (but not full sequence information) is available, we used ePCR [66] with options –g1 –n2 –d50−500 and keeping all successful mappings, except where noted. For cases where locus sequence information is available, we used blastn with an e-value cutoff of 10-8 and retain the top 8 hits for each marker, except where noted. In both cases, we only retain positions where the sequence chromosome and the genetic map chromosome are identical. Specific curation and data cleaning steps for individual species are summarized in Supplemental Table S12 and described in more detail in Supplemental Text S1.

### Removal of incorrectly ordered or duplicated markers

For most species, the genetic position and physical position of markers along a chromosome are not completely congruently ordered. That is, physical position is typically not strictly monotonically increasingly with genetic position. Incongruent markers can arise from incorrect genome assemblies, errors in map construction, or sequence rearrangements between the reference genome and the mapping population.

For consistency, we assume that the reference genome is correctly assembled, and we correct the order and orientation of genetic maps to be consistent with the sequence assembly. To remove incongruent markers, we find the longest common subsequence (LCS) of ranked genetic and physical positions, and define as incongruent all markers that are not part of the LCS. After removing incongruent markers, we filtered each map to retain only the single most congruent mapping position for markers with multiple possible genomic locations. Functions to perform this analysis in R are available at the GitHub page associated with this manuscript.

### Masking low quality map regions

To improve the quality of our recombination rate estimation, we designed a masking filter to exclude regions of chromosomes where the fit between the genetic map and the physical position of markers is particularly poor, defined as a run of 5 bad markers (for chromosomes with at least 25 markers), or a run of 0.2 times the number of markers on the chromosome, rounded up, bad markers (for chromosomes with at fewer than 25 markers). We also completely mask any chromosome with fewer than 5 markers in total. The final map quality and various filtering results are summarized in Supplemental Table S13.

### Recombination rate estimation

Our basic approach to recombination rate estimation is to fit a continuous function to the Marey map relating genetic position and physical position for each chromosome. We use two different approaches that result in different degrees of smoothing: a polynomial fit and a linear B-spline fit. In both cases, we start by optimizing the polynomial degree or spline degrees of freedom using a custom R function that maximizes the Akaike Information Criteria (AIC) for the model fit. For the polynomial fit, we optimize between degree 1 and degree max(3, min(20, # markers / 3)). For the B-spline fit, we optimize degrees of freedom between df 1 and min(100, max(2, #markers/2)). In each case, we retain the value with the highest AIC. To compute recombination rates in cM/Mb, we then take the derivative of the fitted function, evaluated at the midpoint of each window. For additional smoothing, we set all values of recombination estimated below 0 to 0, and all values above the 99th percentile to the 99th percentile. While the two estimates tend to be highly correlated with each other, the polynomial fit appears to perform better for low quality maps, and the B-spline fit for high quality maps. Therefore, unless otherwise noted, we use the polynomial estimates of recombination rate for maps with intermarker spacing of greater than 2 cM, and the B-spline estimates for maps with inter-marker spacing less than or equal to 2 cM. All estimation was done in R; code is available at the GitHub page associated with this manuscript.

### Partial correlations between recombination rate and genetic diversity

To estimate the strength of the association between recombination rate and genetic diversity, we use partial correlations that account for variation in coding sequence density across the genome. In many species [26, 40] recombination rate and/or neutral diversity is correlated with gene density, and thus we need to account for this confounding variable in our analysis. We do this using partial correlations, implemented with the ppcor package in R.

First, we estimate coding sequence density in each window as the fraction of each window represented by CDS sites, extracted from the same GFF files for each species used to compute fourfold degenerate sites. We then estimate Kendall’s τ between recombination rate and genetic diversity for each window after correcting for coding sequence density.

## 4. Modeling the joint effects of background selection and hitchhiking on neutral diversity

We begin with the very general selective sweep model derived by Coop and Ralph [41], which captures a broad variety of hitchhiking dynamics. To include the effects of background selection, we rely on the fact that to a first approximation, background selection can be thought of as reducing the effective population size and therefore increasing the rate of coalescence. This effect can be incorporated by a relatively simple modification to equation 16 of [41]. Specifically, we scale N by a BGS parameter, exp(-G), in equation 16, which then leads to a new expectation of average pairwise genetic diversity (π):

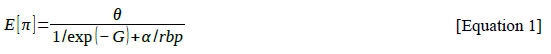

where α = 2N * Vb_p_ * J2,2 (per [41]) and rbp is the recombination rate per base pair. This is very similar to previously published models of the joint effects of background selection and hitchhiking (e.g. [39]). To account for variation in the density of targets of selection, we build upon the approach of Rockman *et al.* [40] and Flowers *et al.* [26], which derives from the work Hudson, Kaplan, Charlesworth and others that originally described models of background selection in recombining genomes [17,18]. Specifically, we fit the following model to estimate G for each window i:

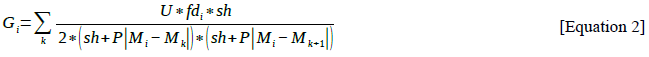

where U is the total genomic deleterious mutation rate, fd_i_ is the functional density of window i, sh is a compound parameter capturing both dominance and the strength of selection against deleterious mutations, M_k_ and M_i_ are the genetic positions in Morgans of window k and window i respectively, and P is the index of panmixis, which allows us to account for the effects of selfing. We estimate functional density as the fraction of exonic coding sites in the genome that fall within the window in question. We focus on exonic coding sites as a proxy for targets of selection as they are the only functional measure that is uniformly available for all the species in our study.

Because P, U, and sh are not known, we fit this BGS model with a variety of parameter combinations. As U is generally unknown, and estimating U is difficult in most cases (*e.g.* [67,68]), we fit our models with three different values: Umin, where we assume U is equal to the mutation rate times the number of exonic protein-coding bases in the genome; Uconst, where we assume that U is equal to 1 for all species; and Umax where we assume that U is equal the lesser of the mutation rate times 5 times the number of exonic protein-coding bases in the genome or the mutation rate times the genome size. Umin and Umax are multiplied by 2 to convert to diploid estimates. We believe that these estimates of U should roughly span the reasonable range for most species. Umin is likely to underestimate the true deleterious mutation rate as the number of exonic protein-coding bases will typically underestimate the number of evolutionarily conserved bases in a genome. Umax assumes that 20% of conserved bases are exonic coding bases and 80% are non-coding, which we admit is a relatively arbitrary assumption, but likely close to the maximum plausible U.

For P, we assume 1 for all vertebrates, insects, and obligate outcrossers among plants; 0.04 for highly selfing species, and 0.68 for partial selfers. These estimates correspond to selfing rates of 0%, ~98%, and ~50%, respectively. Estimates of selfing are available in Supplemental Table S14. For a few species of plants we were unable to obtain reliable estimates of selfing rate (indicated by NA in Supplemental Table S14), and in this case we include all estimates of P in our model selection approach below. For sh, we fit a range of values evenly spaced (on a log scale) between 1e-5 and 0.1. Code to estimate Gi was implemented in C++ and is available from the GitHub repository associated with this manuscript.

To incorporate functional density into the hitchhiking component of the model, we make the simplifying assumption that sweeps targeting selected sites outside a window will have little effect on neutral diversity within a window, and that sweeps occur uniformly within a window. Under this assumption, we can consider functional density as a scaling factor on the rate of sweeps, Vbp. Specifically, we reparameterize the rate of sweeps, Vbp, as V, the total sweeps per genome, and then consider the fraction of sweeps that occur in a particular window i as V*fd(i). This results in a simple scaling of alpha in equation 1. While we note that this assumption is likely to be violated in practice, it allows use to use the homogeneous sweep model of [41] with different rates of sweeps for each window across the genome. Ultimately, of course, it would be preferable to derive a non-homogenous sweep model that directly incorporates variation in functional density, but doing so is beyond the scope of this manuscript. However, we believe that our simplifying is likely adequate, as the largest reduction in diversity associated with a sweep is localized to the window containing the swept site (*e.g.,* [41]).

Incorporating the effects of functional density in both BGS and HH, our final model for the expectation of neutral diversity in window i is:

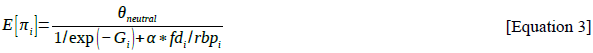

To obtain an estimate of the effect of selection for each species, we fit this model for estimates of G_i_ derived from different parameter combinations (see above), using the nlsLM() function from the minpack.lm package in R. In addition, we fit three simpler models: a background selection only model (in which alpha is 0 and thus the second part of the denominator is null), a hitchhiking only model (in which G is 0 for all i, and thus the first part of the denominator is null), and a neutral model in which both G and alpha are 0, and thus the model predicts that neutral genetic diversity is equal to mean genetic diversity across the genome. Together, we refer to these four models as model set 1. Finally, we fit a second set of models (model set 2) in which we use the same approach to model background selection, but use homogenous hitch-hiking model of [41] without modification to allow for variation in functional density across the genome, and thus remove the fdi term from (Equation 3).

From each model fit we estimate theta[neutral] for all four models (full, BGS-only, HH-only, neutral), and also extract the likelihood of the fit. We then compute the AIC for each parameter combination, extract the fit with the best AIC for each model, and use the AIC to estimate the Akaike weight (relative likelihood) of each model j as

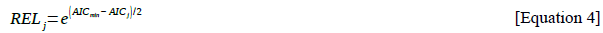

which we then normalize so that the weights for all four models within a for a species sum to 1. We focus on AIC as it provides a straightforward way to compare non-nested models.

We estimate neutral genetic diversity for each species as the parameter value obtained by the model with the best AIC. We then compute average observed genetic diversity for each species, and report the magnitude of the impact of selection on linked neutral diversity as 1 – (observed/neutral). Values below 0 are replaced by 0. This value can be interpreted as the proportion of neutral variationremoved by selection acting on linked sites, averaged across the genome.

This modeling approach has some important limitations: in particular, our approach calculates the effects of BGS and HH in windows across the genome instead of per base and we use the parameter *sh* instead of integrating across the distribution of fitness effects (as is done in *e.g.* [48,50]). Additionally, we do not use information such as locations of amino acid fixations, as is used by *e.g.* [49]. We fully acknowledge that these simplifying assumptions will, to a certain extent, degrade the accuracy of our modeling approach compared to other possible approaches. We argue, however, that these assumptions are necessary for this work: more sophisticated models typically require additional data (e.g., the distribution of fitness effects of new mutations or the location of recent amino acid fixations), or significantly increased computational time (i.e., by computing the effects of background selection at each base instead of in windows). For most of the species we studied, the necessary additional data are not clearly available to fit more complex models, and the increased computational time to per-base models would rapidly make our analysis computationally intractable. Thus, we believe that we have made reasonable tradeoffs between modeling complexity, data availability and taxonomic breadth.

## 1. Linear models

Our goal is to test whether Nc predicts the degree to which selection shapes patterns of neutral diversity, using log-transformed measures of body size and geographic range as proxies for Nc. However, many other factors could potentially influence our measure of strength of selection, including biological factors such as genome size and average recombination rate; and experimental factors such as map quality and assembly quality. In particular, we might expect to underestimate the strength of selection in species with low-quality assemblies or maps, and we might expect that on average larger genomes and higher recombination rates would reduce the impact of selection.

In order to account for these parameters that are not directly of interest, we use two approaches. First, we compare a model that includes both our parameters of interest and our parameters not directly of interest to a model that includes only the parameters not directly of interest, in order to test whether our proxies for Nc result in a significantly better fit. Second, we fit our proxies for Nc to the residuals of a linear model including only parameters not directly of interest, in order to determine how much variation proxies for Nc explain after accounting for all the variation that can be explained by genome size, average recombination rate, and quality parameters.

We obtain assembly quality from NCBI, Phytozome, the original genome publication, or compute it directly from fasta files. C-values for plants come from http://data.kew.org/cvalues/, and C-values for animals come from http://www.genomesize.com/. In all cases, most recent estimates, “prime” estimates, or flow cytometry estimates are preferred; where several seemingly equally good estimates are available, the average is used. In some rare cases a related species is used instead of the sequenced species if the sequenced C-value is not available. We focus on C-values instead of assembly size as using assembly size as a measure of genome size confounds genome size and assembly quality (lower quality assemblies will be on average less complete and therefore smaller). Assembly parameters and sources are listed in Supplemental Table S15. Average recombination rate computed as the overall map size divided by the size of the genome covered by the map.

In order to determine which interactions among proxies for Nc (size, range, and kingdom) to include, we start with the full model including all interactions and remove all non-significant interactions. After doing so, we our model is

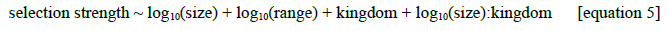

## 6. **Data accessibility**

The data we analyze in this manuscript, and the scripts we used to produce our results, are available as follows. All genomes, polymorphism datasets, and GFF annotation files are publicly available from NCBI or other sources. Genome references and versions are listed in Table S1, and URLs pointing to the location of genome sequence and GFF annotations are available in Table S2. SRA accessions for polymorphism datasets are listed in Table S10, and references for polymorphism datasets, where available, are listed in Table S1. Genetic maps for each species are available from the references listed in Table S1, or as an R data file available at the GitHub page associated with this manuscript (https://github.com/tsackton/linked-selection). All Perl scripts, R scripts, and C++ code associated with this manuscript is available from GitHub (https://github.com/tsackton/linked-selection). and the function of each piece of code is documented both in comments in the code itself and in the Github README. Programs used for read mapping and genotyping, along with command line parameters, are described in the methods. The GitHub page also provides several intermediate data files, including range and size data for each species, neutral diversity and recombination rate for 100kb, 500kb, and 1000kb windows across each species, and the final dataset analyzed with the linear models described above.

## Acknowledgements

The authors acknowledge helpful comments and advice from Daven Presgraves, Joshua Schraiber, Rasmus Nielsen, Chuck Langley, John Wakeley, Graham Coop, Andrew Berry, Andrew Clark, Julien Ayroles, Shelbi Russell, Brian Arnold, and Jacob Crawford, as well as three anonymous reviewers whose comments greatly improved the manuscript. The computations in this paper were run on the Odyssey cluster supported by the FAS Division of Science, Research Computing Group at Harvard University.

## Author contributions

RBC-D and TBS conceived and designed this study, processed and analyzed the data, and wrote the manuscript. DLH offered advice and contributed to writing the manuscript. All authors read and approved the final version.

## Supplemental materials

Table S1: details of species used

Table S2: links to online data sources for species used

Table S3: correlation robustness analysis

Table S4: model fit to residuals of nuisance parameter model

Table S5: model fit to residuals of nuisance parameter model (minus genome size)

Table S6: model fit after removing domesticated species

Table S7: model fit to animals only

Table S8: model fit to plants only

Table S9: list of sizes for each species along with source

Table S10: accessions for polymorphism data

Table S11: sites aligned for each species

Table S12: summary of map curation / cleaning steps for each species

Table S13: final map quality summary

Table S14: estimates of selfing and associated references

Table S15: quality parameters used in linear modeling

Supplemental Figures:

Figure S1: bioinformatic pipeline schematic

Supplemental Text:

Text S1: details of map construction for each species Supplemental References: references from supplemental tables

